# Psilocybin increases optimistic engagement over time: computational modelling of behavior in rats

**DOI:** 10.1101/2024.05.16.594614

**Authors:** Elizabeth L. Fisher, Ryan Smith, Andrew W. Corcoran, Laura K. Milton, Kyna Conn, Jakob Hohwy, Claire J. Foldi

## Abstract

Psilocybin has shown promise as a novel pharmacological intervention for treatment of depression, where post-acute effects of psilocybin treatment have been associated with increased positive mood and decreased pessimism. Although psilocybin is proving to be effective in clinical trials for treatment of psychiatric disorders, the information processing mechanisms affected by psilocybin are not well understood. Here, we fit computational models of underlying decision-making mechanisms to behaviour in rats. The model revealed that rats treated with psilocybin achieve more rewards through increased task engagement, mediated by modification of forgetting rates and reduced loss aversion. These findings suggest that psilocybin may afford an optimism bias that arises through altered belief updating, with translational potential for clinical populations characterised by lack of optimism.

## Introduction

Psilocybin has shown promise as a novel pharmacological intervention for treatment of depression, where post-acute effects of psilocybin treatment have been associated with increased positive mood and decreased pessimism (1–3). Although accumulating evidence indicates that psilocybin is effective for treatment of psychiatric disorders, the information processing mechanisms underlying the effects of psilocybin are not well understood (4,5). Establishing the information processing mechanisms of psilocybin could significantly benefit our understanding of the drug’s therapeutic actions, potentially helping to improve its efficacy and specificity, which could assist in informing clinical decisions. One way we can investigate the post-acute effects of psilocybin is with animal models (6,7), which has a number of benefits, including the ability to collect many data points in controlled environments and over sustained periods after treatment, as well as circumventing issues with expectancy effects that may confound the results from clinical trials in humans (8).

To understand the information processing mechanisms of psilocybin, computational modeling approaches allow us to investigate the change in specific model parameters over time, providing insight into how psilocybin may help treat depression. Such computational modeling approaches fall into the burgeoning field of computational psychiatry that aims to develop precise treatments for psychiatric disorders, based on the specific information processing mechanisms of a particular individual, with an understanding that shared symptoms may arise from different computational processes (9–13). Here, we employed a novel two-armed bandit reversal learning task capable of capturing engagement behaviour in rats. Measuring engagement behaviour has translational potential for individuals with depression, who often choose to withdraw from the world rather than engage in rewarding activities (14,15). In fact, modifying such behaviour is a primary target of existing behavioural activation interventions within cognitive-behavioural therapies (16,17). In our experiment, two groups of rats were administered psilocybin (n=12) or saline (n=10) 24 hours before the initiation of the task. The rats could engage with the task for three hours each day for 14 days and completed the task in their home cage such that they could decide to either engage in the task or stay in the cage without engaging.

The present study used both behavioural measures and computational modelling methods to distinguish the mechanisms underlying task engagement. The space of models included both reinforcement learning (RL) (18) and active inference (AI) (19–21) models, with a range of possible parameters motivated by prior research on depression. As optimism bias is associated with increased engagement with the world and depression is linked to diminished optimism, we include parameters related to optimism, allowing us to investigate if increased engagement in the task can be account for by increased optimism (22–26). We hypothesized that psilocybin would increase subsequent task engagement through discrete changes in information processing in rats.

## Methods

### Animals

All animals were obtained from the Monash Animal Research Platform. Female Sprague-Dawley rats (n=22) arrived at 6 weeks of age and were allowed to acclimate to the reverse light cycle (lights off at 11AM) for 7 days prior to any intervention. Young female rats were used in these studies to compare outcomes to previous studies in the laboratory investigating the role of psilocybin on cognitive flexibility (27). Because reinforcement learning motivation is known to fluctuate over the estrous cycle (28), a male rat was housed in all experimental rooms at least 7 days prior to experimentation, synchronizing cycling (cf. the Whitten Effect (29)). Before training, rats were housed individually in specialized cages (26cm W x 21cm H 47.5cm D) and were habituated to sucrose pellets (20 mg, AS5TUT, CA, USA) sprinkled into the cage for two consecutive days. Pre-training involved first training animals to take pellets from the magazine (magazine training) on a “free-feeding” schedule, and subsequently training animals to make nose-poke responses to obtain a pellet (nose-poke training), a nose-poke into either port delivered a pellet at a fixed ratio (FR) 1 schedule (See Supplementary Figure 1). At the completion of pre-training, when rats were between 8-9 weeks of age, a single dose of psilocybin (1.5 mg/kg; USONA institute, dissolved in saline) or saline alone (control) was administered intraperitoneally and reversal learning training commenced the following day. Psilocybin efficacy was confirmed by the adoption of hind limb abduction, a stereotypical posture. All experimental procedures were conducted in accordance with the Australian Code for the care and use of animals for scientific purposes and approved by the Monash Animal Resource Platform Ethics Committee (ERM 29143).

### Reversal Learning Task

For the reversal learning task, Feeding Experiment Devices (version 3; FED3) (30) were placed in the home cage of the rat for 3 hours a day (**Figure 1**). The rats were maintained on *ad libitum* access to food (standard rat and mouse chow; Barastoc, Australia) throughout the entire experimental paradigm in order to rule out hunger as a motivating factor for performance. The reversal learning task was a modified two-arm bandit designed so that 10 pokes on the rewarded side of the FED3, with each poke resulting in a reward, triggered the reversal of the rewarded side. This continued on a deterministic schedule of reinforcement (i.e., “active” pokes delivered a reward 100% of the time) over 14 experimental sessions on separate and consecutive days, where the left side was active first. There were three outcomes from the rats’ actions in the reversal learning task. If the rat poked the active side and received a sucrose pellet, the outcome was a ‘reward’. If the rat poked the non-active side of the FED3 the outcome was a ‘loss’, as they exerted energy without reward. If the rat did not engage in the task, the outcome was ‘null’, which was considered a better outcome than a loss as the rat did not exert energy without reward.

**Figure 1.**
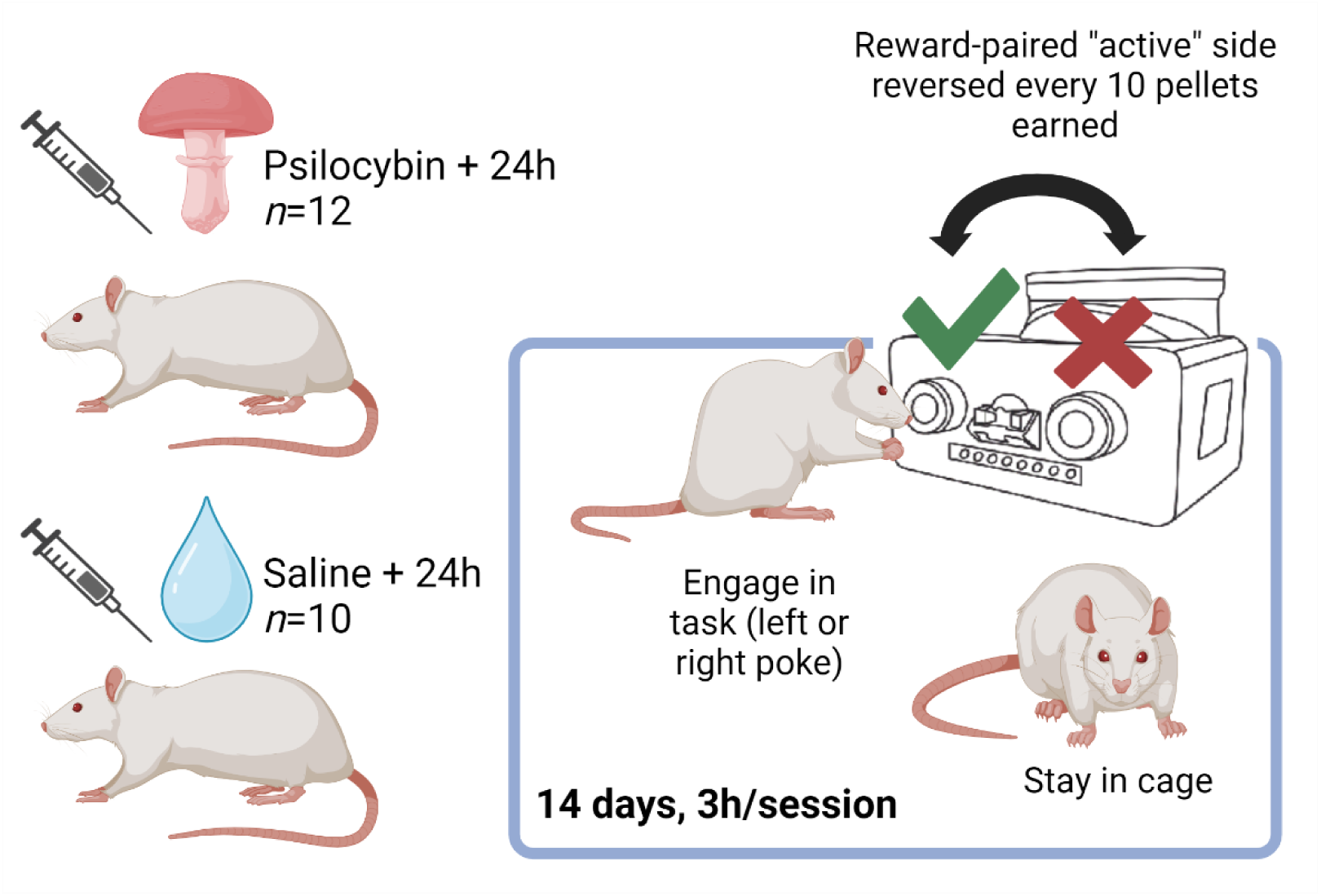
Treatment and reversal learning task design. Rats completed a reversal learning task over 14 experimental sessions on separate consecutive days in their cage, with the first session beginning 24 hours after treatment with psilocybin (n=12) or saline (n=10). The reversal learning task had a deterministic reinforcement schedule and was designed so that 10 pokes on the rewarded side, with each poke resulting in a reward, triggered the reversal of the rewarded side. The rat could select from three actions: poke left, poke right, or stay in the cage. Created with biorender.com

### Locomotor activity and anxiety-like behavior

To examine potential influences of general locomotor and anxiety-like behavior that may contribute to task performance, a separate cohort of female Sprague-Dawley rats (age-matched to 8-9 weeks old) were administered psilocybin (*n*=8) or saline (*n*=8) and 24h later were tested in the open field (OF) and elevated plus maze (EPM). Behavioral tests were recorded with an overhead camera connected to a computer and analyzed with Ethovision XT (V3.0; Noldus, NL) tracking software using center of mass. The EPM consisted of an elevated 4-arm platform made of grey Perspex (70 cm long × 10 cm wide × 90 cm high) with two closed (40 cm high walls) and two open arms. Rats were placed in the centre platform (10 × 10 cm) facing an open arm and the proportion of time spent in the closed arms relative to the open arms in each 10-min trial, was used as the primary measure of anxiety-like behaviour (31). The OF test consisted of a deep open topped box (60 x 60 x 55 cm deep) in which distance travelled in each 10-min trial was used as the primary measure of locomotor activity and the proportion of time spent in the aversive centre zone (middle square of a 3 x 3 grid; 20 × 20 cm) was used as a secondary measure of anxiety-related behaviour, although it is shown to be less sensitive to the effects of anxiolytic drugs (32).

### Computational Modelling methods

We considered reinforcement learning (RL) (18) and active inference (AI) (19–21,33) models as possible explanations of observed choice behavior. For all computational models, the rat had three actions to choose from: ‘poke left’, ‘poke right’, or ‘stay in cage’, and three outcomes: ‘reward’, ‘loss’, or ‘null’. A reward was modelled as a positive outcome, loss as a negative outcome, and null as a neutral outcome (i.e., less aversive than a loss, as the rat conserved energy).

#### Reinforcement Learning models

Four different RL models were considered that reflect hypotheses about how optimism bias might be computationally implemented. We hypothesized psilocybin increased optimism by asymmetric belief updating, which was the main focus of these models; however, other parameters that have been associated with depression or optimism were also considered. Note that, we only include the most common RL modeling approaches here that do not include explicit beliefs about state transition probabilities (i.e., so-called “model-free” RL). However, more complex models with explicit transition beliefs (i.e., “model-based” RL) could also be considered (33).

All models included, *V*_0_, which is the initial value of the expected reward for each action, ‘stay in cage’, ‘poke left’, ‘poke right’. The initial value of the expected reward was restricted to positive values. The value of the expected reward was transformed into a discrete probability distribution using a softmax function, which included a standard inverse temperature parameter (*β*) controlling randomness (value insensitivity) in choice, as follows where *a* is set of possible actions.

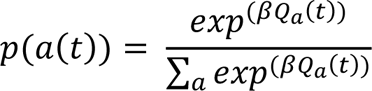

Here, trial is denoted by *t*. Expected values (*Q*) were updated based on reward prediction errors (*RPE*). These prediction errors reflect the difference between the current (*r*) and the expected reward, as follows:

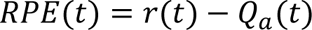

The reward value was coded as 1 for a reward, -1 for a loss, and 0 for the null outcome.

##### Simple Rescorla-Wagner

The simplest RL (Rescorla-Wagner) model included one learning rate parameter, *α*, in addition to *V*_0_ and *β*, The expected reward was here updated at the same rate regardless of whether the rat observed a reward or a loss.

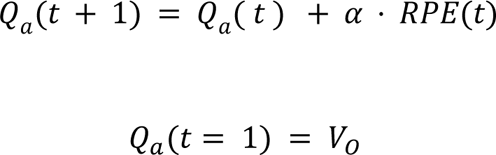

##### Pessimistic Rescorla-Wagner

The pessimistic Rescorla-Wagner model aimed to account for a pessimism bias the rat might possess. For this model, we fixed the initial value for the expected reward for the ‘stay in cage’ action to be 0. During fitting, the estimated initial values of *V*_0_ for the ‘poke left’ and ‘poke right’ actions were then permitted to take negative or positive values. This allowed the rat to potentially begin the trial with a pessimism bias, in which it initially believed that selecting left or right sides would lead to a negative outcome (promoting avoidance), which would need to be unlearned through subsequent task engagement.

##### Asymmetric Learning model Loss Aversion

As depression and optimism has been associated with differences in belief updating to good versus bad outcomes (34), we tested a Rescorla-Wagner model with separate learning rates for rewards and losses.

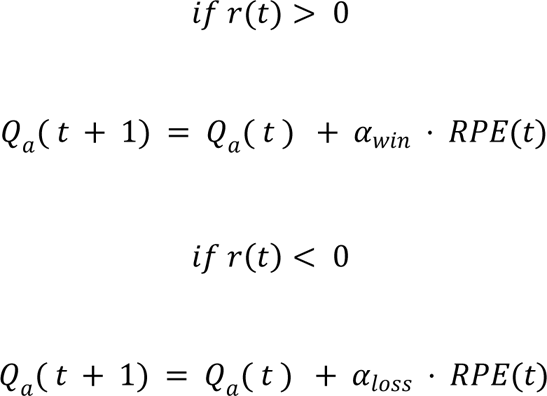

Additionally, as depression is related to increased loss aversion, this model included a loss aversion parameter (22). This parameter was implemented as the negative reward value assigned to the loss outcome. Instead of being encoded as a -1 (as in the other models above), this negative reward could take larger values if a rat was more loss averse. Thus, under a loss outcome, the negative reward was scaled by a loss aversion (LA) parameter as follows:

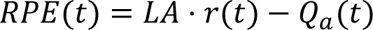

##### Dynamic Learning Rate (Associability)

Theoretically, learning rate should be greater when the environment is expected to be volatile/unpredictable, while learning rate should decrease in a stable/predictable environment. Prior work suggests this learning rate adjustment may be altered in affective disorders (35). To evaluate the possibility that differences in learning rate adjustment could account for differences in optimistic behavior, we included an ‘associability’ RL model that allows learning rate to be adjusted on a trial-by-trial basis, based on the magnitude of previous prediction errors (i.e., larger prediction errors effectively increase learning rates, as they suggest confidence in current expectations should be low). This model includes a parameter (η), which modulates how strongly the strength of learning rate can change after each trial. The update equations for associability (κ), which is used to modulate the baseline learning rate on each trial, are as follows:

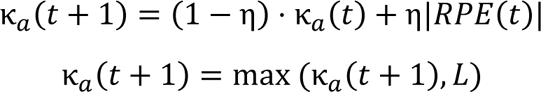

Note that a lower bound (*L*) on the value of κ is required, which we fit as an additional parameter. The expected reward is then calculated by the following equation:

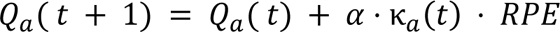

### Active Inference models

In addition to RL models, we consider active inference (AI) models, which incorporate Bayesian learning and an intrinsic information-seeking drive that could play a role in promoting engagement behavior (**Figure 2**). The categories of *actions* and outcomes (*o*) were set in the same manner here as in the RL models. The main differences from RL, as described further below, are that: 1) the model explicitly learns probabilities of each outcome under each action and estimates its confidence in current beliefs about those probabilities; 2) the reward value of each outcome is encoded in the form of a probability distribution, *p*(*o*|*C*), where higher probabilities correspond to greater subjective reward; and 3) action selection is driven by an objective function called expected free energy (*Q*), which jointly maximizes expected reward and expected reductions in uncertainty about outcome probabilities.

**Figure 2.**
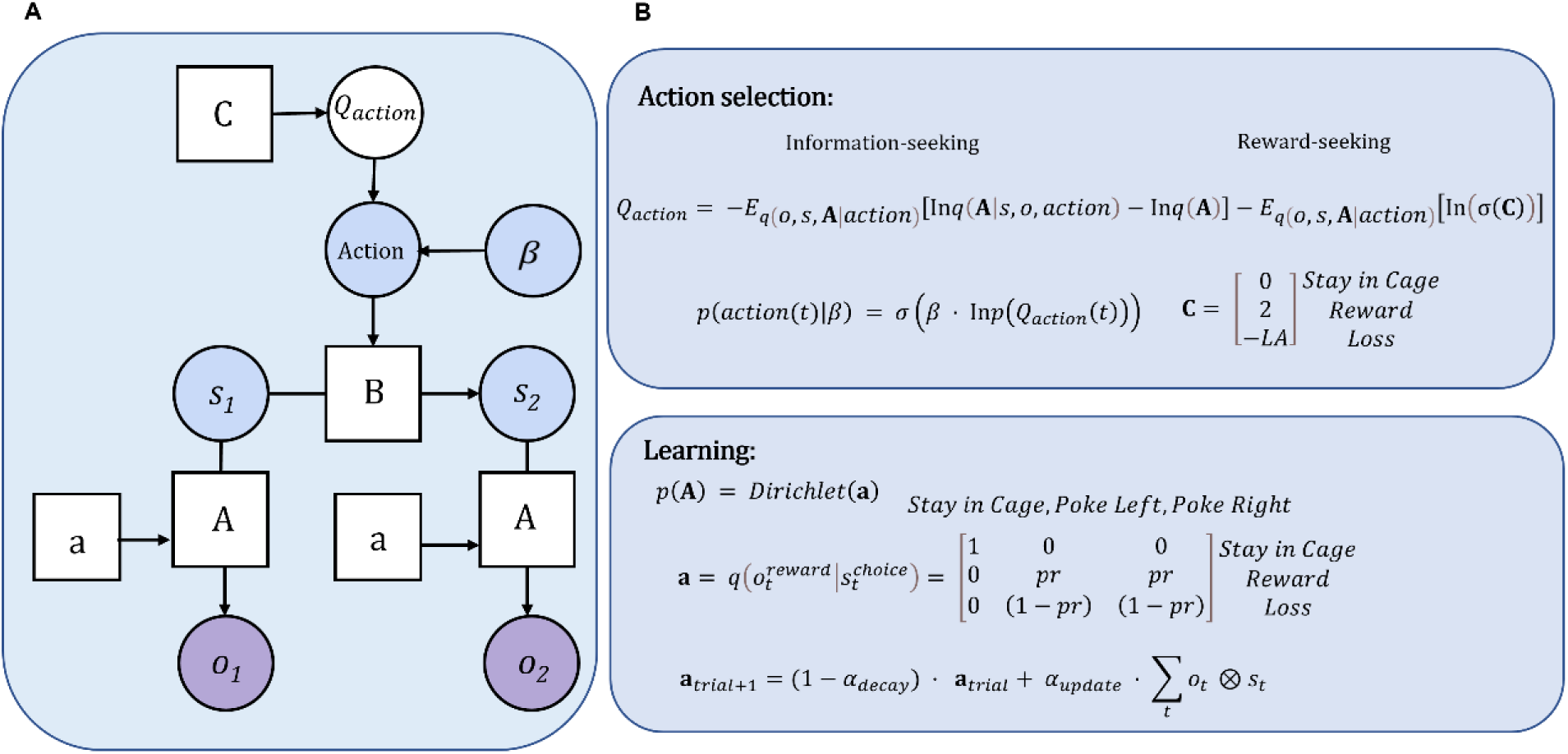
Active inference model. **2A:** A graphical depiction of the active inference model. **2B:** Active inference model equations for learning and action selection. **C** is the subjective reward value for each outcome. *Q*_*action*_ is the action value for each *action*. *s* is the action states. **B** is the state transition probabilities. **A** is the reward probabilities of states and outcomes in the task and *a* is the prior beliefs about the reward probabilities. *o* is the action observations. The *σ* symbol indicates a softmax function. The right two panels show the model equations for action selection and learning. See *Methods* for more details on the model and equations.

Expected free energy is calculated by the following equation:

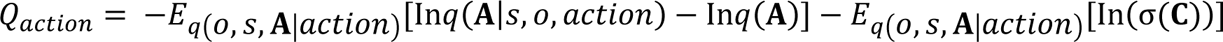

Here, **A** is a matrix that encodes the reward probabilities, *p*(*o*|*s*), as the relationship between action states (*s*) and observations (*o*). **C** is a vector that encodes the reward values for the rat. The *σ* symbol indicates a softmax function. The first term in the expected free energy equation calculates how much information about the reward probabilities would be gained by taking an action. In other words, if the rat selects a certain action, the magnitude of this term indicates how much it expects to increase its confidence in the best action. In this model actions are considered observable. Note here that the symbol *q* indicates an approximate probability distribution. The second term in *Q*_*action*_ calculates the expected probability of reward under each action, based on current beliefs.

The first AI model included five parameters with theoretical links to depression and optimism, similar to those in the RL models. To investigate the differences in belief updating from positive and negative outcomes associated with depression and optimism (34), we included separate forgetting rate parameters (*α*_*decay*_) for rewards and losses. In the AI framework, forgetting rates are conceptually similar to (but mathematically distinct from) learning rates in RL, as they indicate how strongly the rat forgets its prior beliefs before observing a new reward or loss. The higher the value for the forgetting rate, the more the rat updates its beliefs after a new observation (i.e., larger values down-weight prior beliefs before each update). Learning specifically involves updating the parameters of a Dirichlet distribution (**a**) over the likelihood matrix **A** after each observation, as shown in the equation below. We included a fixed value (0.5) for learning rate (*α*_**u***pdate*_).

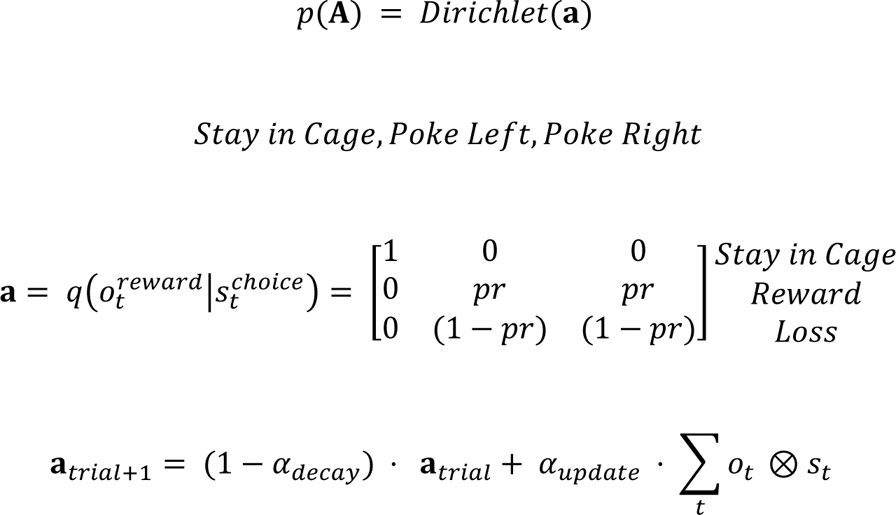

We included an initial prior value for probability of receiving a positive outcome as an additional parameter (*pr*) (i.e., the initial expected probability of a reward under both actions at the start of the task, prior to learning). As optimism is associated with expectations of a good outcome (26,36).

Similar to the RL model space, we included a loss aversion parameter (*LA*) encoding the value of a loss, assigning a fixed positive value of 2 for observing reward:

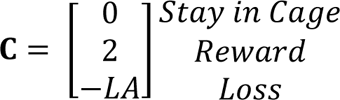

We included an inverse temperature (action precision) parameter (*β*) that controlled the level of randomness in action selection:

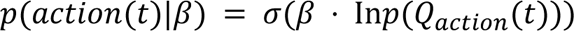

Finally, we compared this model to a simpler four-parameter AI model that omitted the prior reward probability (*pr*) parameter, where *pr* is fixed at 0.5.

#### Model comparison

Model simulations for the AI and RL models were run in MATLAB (v.9.12.0, (R2022a)). To estimate the parameters for all models, a variational Bayes algorithm (variational Laplace) was used, which maximizes the likelihood of the rat’s actions while incorporating a complexity cost to deter overfitting (37).To find the best model, we then used Bayesian model comparison across the 4 RL models and 2 AI models (38). To confirm the recoverability of parameters in the best model, we then simulated behavioral data using the parameters we estimated for each rat (based on the same number of trials each rat performed in the task). Using this simulated behavioral data, we then estimated these parameters and evaluated how strongly the estimates correlated with the true generative values. Recoverability was acceptable for all parameters in the winning model (see **Supplementary Materials**).

### Statistical analysis

All analyses were performed in R (v4.1.1) using functions from the Tidyverse package (v1.3.2; (39)). To explore whether model parameters might jointly differentiate groups, we performed logistic regression analyses with the parameters as predictors and group classification as the outcome variable. Interactions between each parameter and time (day) were included. This was carried out using the glm() function in R with the ‘binomial’ family and the ‘logit’ link function.

Behavioral performance and computational model parameters were modelled using generalized additive mixed-effects models (GAMMs), a semiparametric extension of the generalized linear mixed-effects modelling framework that enables the fitting of nonparametric curves (splines) describing how the mean of an outcome variable *y* varies as a smooth function *f* of explanatory variable(s) *x* (40–42). A smoothing penalty is applied during the fitting of smooth terms, ensuring that the complexity (i.e., ‘wiggliness’) of *f*(*x*) is appropriately constrained (e.g., reducing *f*(*x*) to a linear function in the event there is insufficient evidence of nonlinearity in the mapping between *y* and *x*).

GAMMs were estimated using restricted maximum likelihood via the gam() function of the mgcv package (v1.8-40; (42)). All GAMMs took the following generic form:

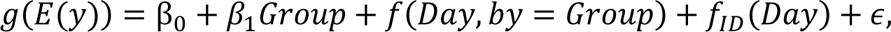

where Group encoded a two-level factor variable reflecting treatment allocation (psilocybin, control) and Day encoded the consecutive sequence of test sessions (1–14). Thin-plate regression splines (43) were used to model changes in *y* over Day for each level of Group. Factor smooths on Day were implemented for each individual rat (ID) to capture random fluctuations in behaviour over successive sessions (this term essentially functions as a time-varying random intercept; for similar approaches, see Corcoran et al. (44); Cross et al. (45)). The selection of link function *g*(.) was informed by the distribution of *y* and quality of model fit, which was evaluated using functions from the mgcViz (v0.1.9; (46)) and itsadug (v2.4.1; (47)) packages.

To understand the relationship between the behavioral outcomes and the winning model parameters, we ran Pearson correlations between each model parameter and behavioral outcome measure.

## Results

### Behavioral Results

Rats treated with psilocybin tended to achieve more rewards in the reversal learning task than those treated with saline (*b*=36.1, *t*(239)=1.92, *p*=.056). There was a significant difference between smoothers; rats in the psilocybin group achieved significantly more rewards on days 7 - 14 compared to those in the control group (**Figure 3A**). Rats in the psilocybin group showed significantly more losses on average than the control group (*b*=15.60, *t*(218)=2.04, *p*=.042). Comparison of smoothers indicated that this difference was driven by fewer losses in the control group on days 4 – 6 (**Figure 3B**). As task engagement was optional, one group could have both more wins and more losses than another group due to more frequent engagement.

**Figure 3.**
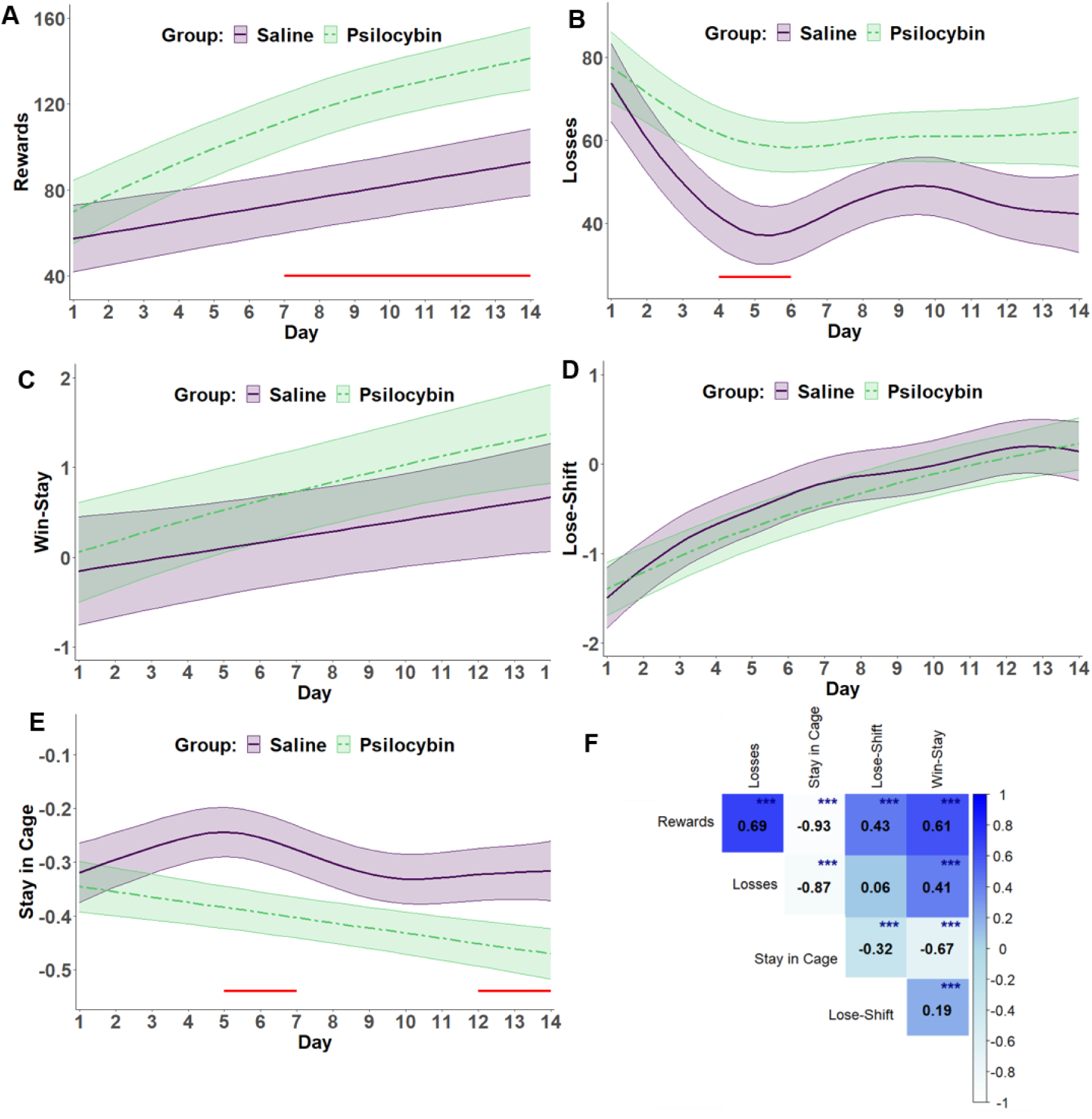
Effects of psilocybin on behavioral measures across days of reversal learning. The first 5 panels in this figure illustrate pairwise comparisons of the behavioral measures across days including (A) rewards, (B) losses, (C) win-stay strategies (logit-scaled), (D) lose-shift strategies (logit-scaled), and (E) stay in cage behavior (log-scaled). Estimated marginal means are represented with dotted lines (psilocybin group) or solid lines (saline group) with shading indicating SEM. Solid red lines indicate significant differences between groups (p < .05). Correlations between behavioral measures presented in (F). *p < .05, **p < .01, ***p < .001.

We found no difference in win-stay or lose-shift behavior between the two groups (**Figures 3C, 3D**). In contrast, the control group selected the action of ‘staying in the cage’ significantly more than the psilocybin group (i.e., reflecting less frequent task engagement; *b*=-0.11, *t*(221)=-1.99, *p*=.048). Comparison of smoothing splines further revealed significant reductions in psilocybin group stay-in-cage times on days 5 - 7 and 12 - 14 (**Figure 3E**).

Rewards showed a strong positive association with the amount of time rats chose to engage in the task (*r*=0.93, *p*<.001). Rewards were positively associated with losses (*r*=0.69, *p*<.001). This was because the chances of reward and loss both increased with greater task engagement (**Figure 3F**).

#### Model Comparisons

When comparing four reinforcement learning (RL) and two active inference (AI) models with Bayesian model comparison we found the winning model to be the five parameter AI model (protected exceedance probability = 1). Model validation and parameter recoverability analyses confirmed the validity and reliability of this model (see **Supplementary Materials**).

#### Active Inference Model Results

The inter-correlations for the winning AI model parameters and behavioral results showed several associations in expected directions (**Figure 4F**). The strongest association was with number of rewards received and the loss aversion parameter, where lower loss aversion was correlated with more rewards. Rewards were associated with both forgetting rates in the expected directions.

**Figure 4:**
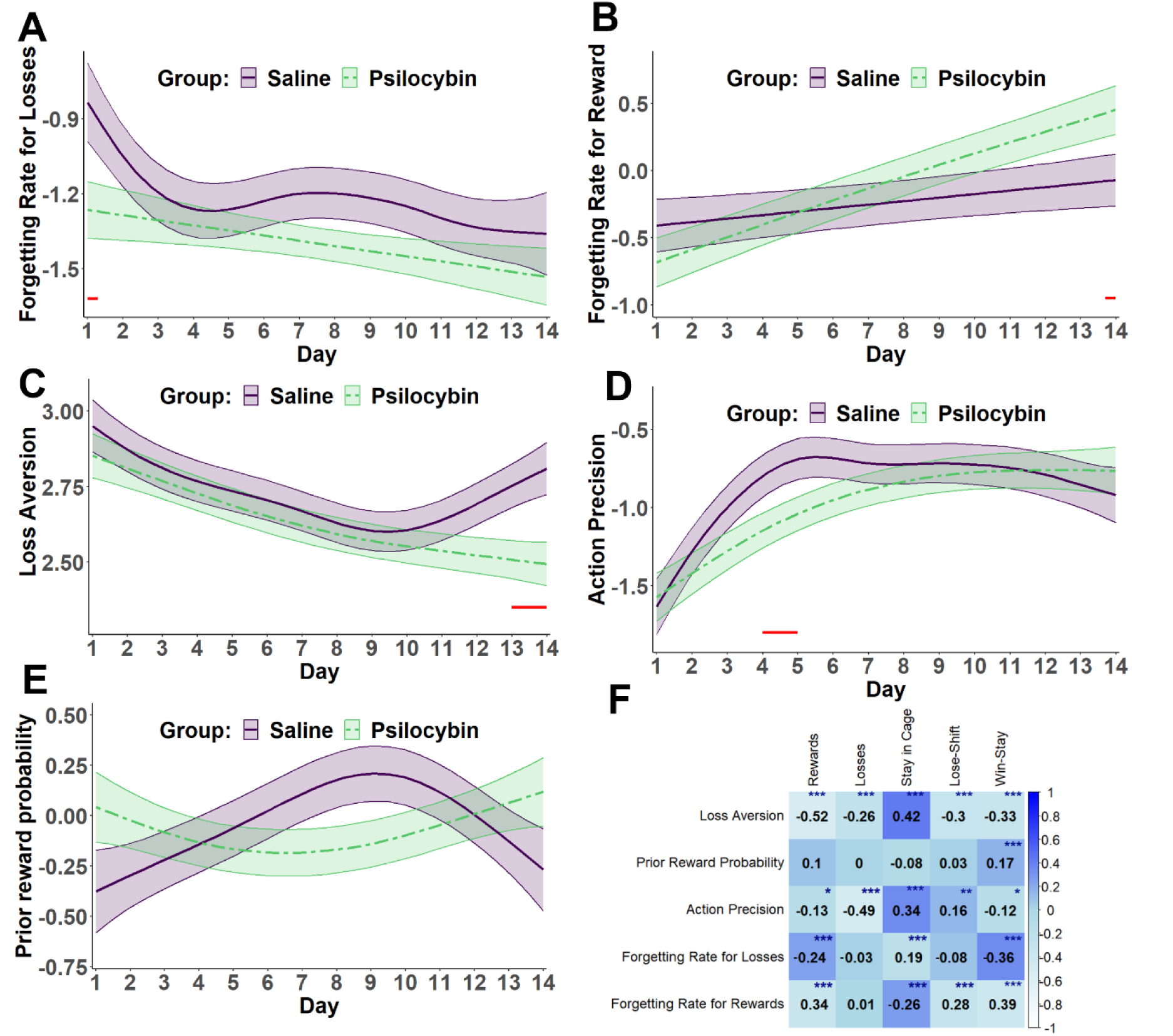
Effects of psilocybin on model parameters across days of reversal learning. The first 5 panels in this figure illustrate pairwise comparisons of the model parameters across days including (A) forgetting rate for losses (logit-scaled), (B) forgetting rate for rewards (logit-scaled), (C) loss aversion (log-scaled), (D) action precision (log-scaled), and (E) prior reward probability (logit-scaled). Estimated marginal means are represented with dotted lines (psilocybin group) or solid lines (saline group) with shading indicating SEM. Solid red lines indicate significant differences between groups (p < .05 Correlations between parameters in the best-fit computational (AI) model and behavioral results measures (F). *p < .05, **p < .01, ***p < .001.

The logistic regression capturing the variance among all parameters found forgetting rate for rewards was significantly higher in the psilocybin group (*b=*.62*, z*(295)=2.034, *p* = .042) and forgetting rate for losses was significantly lower in the psilocybin group (*b=*.57, *z*(295)=-2.053, *p*=.040). We found a significant interaction between forgetting rate for rewards and day (*b=-*.080, *z*(295)=2.294, *p*=.022), indicating that the forgetting rate for rewards for the psilocybin group increased more over time. There was a significant interaction between loss aversion and day (*b=-*.11, *z*(295)=-2.580, *p*=.010), indicating that the psilocybin group had smaller loss aversion on the later days of testing. Taken together, the logistic regression found that parameter space change for psilocybin group compared to saline group was as follows: forgetting rates for rewards were higher, forgetting rates for losses were lower, forgetting rate for rewards increased more over time, and loss aversion was lower for the psilocybin group over time.

Results for the GAMMS, which did not take into account the other parameters in the parameter space, showed forgetting rates for losses were significantly lower on average in the psilocybin group (*b*=0.18, *z*(267)=-1.99, *p*=.046). This between-group difference was most pronounced on day 1, but remained present across the experiment (**Figure 4A**). Although there was no main effect of group for the forgetting rate for rewards, comparison of smoothers revealed this parameter to be significantly higher in the psilocybin group on day 14 (**Figure 4B**).

None of the remaining model parameters showed significant between-group main effects. However, smoothing splines differed significantly between groups for both the loss aversion and action precision parameters on specific days. Loss aversion was lower in the psilocybin group on day 13 and 14, suggesting that this reduced anticipated aversiveness of losses over time (**Figure 4C**). Action precision was significantly lower for the smoothing splines in the psilocybin group on days 4 and 5 (**Figure 4D**). As action precision indicates less randomness in choice, this most likely reflected the fact that rats in this group less deterministically chose to stay in their cage and not perform the task on those days.

#### Other aspects of behavior relevant to task performance

Because engagement during the reversal task could be influenced by prior experience with sucrose rewards, response vigor, general locomotor activity and anxiety-like behavior, we assessed these aspects of behavior separately. During pre-training (**Figure 5A**), where poking into either side of the FED3 resulted in a pellet, there was no effect on psilocybin overall (**Figure 5B**; *p*=.102) and while both groups preferred the right over the left nose-poke (**Figure 5C-D**), initial group differences in sucrose collection did not contribute to performance after treatment with psilocybin or saline. In contrast, we found that during the reversal task (**Figure 5E**), the rate of responding (response vigor) showed a trend toward increase vigor after psilocybin treatment for the target (rewarded) nose-poke over the 14 days (**Figure 5F**; *p*=.096) but this was not the case for non-target (incorrect) responses (**Figure 5E-G**).

**Figure 5:**
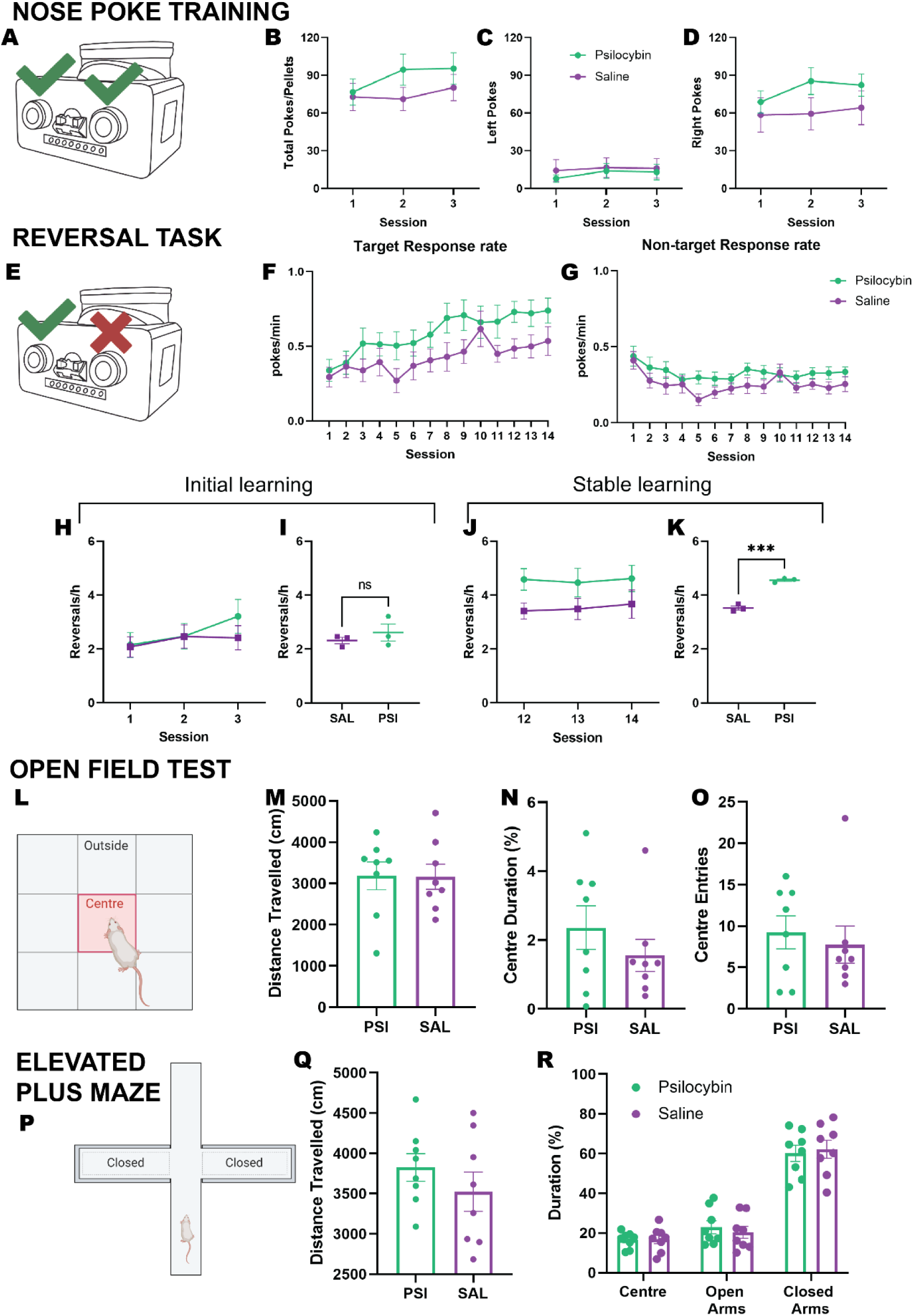
Raw behavioral data. **A-D:** Nose-poke training results prior to psilocybin or saline administration. **E-G:** Response rates for psilocybin and saline groups in the reversal learning task. **H-K:** Reversal rates for psilocybin and saline groups during initial and stable performance periods **L-O:** Open field data for rats treated with psilocybin and saline aged matched to cohort in current study. **P-R:** Elevated plus maze data for rats treated with psilocybin and saline aged matched to cohort in current study. PSI; psilocybin, SAL; saline. Data presented are mean ±SEM.

We found that post-administration day-to-day performance variability, as measured by reversal rate, had stabilized by the late reversal learning period (**Figure 5J**; sessions 12-14, *p*=.681), where significantly higher rates of reversal were observed for the psilocybin treatment group (**Figure 5K**; *p*>.001)This improvement in reversal ability in the later sessions also coincided withthe greatest trend towards an effect of psilocybin on total number of rewards received (*p*=.083) Taken together, this suggests that psilocybin could enhance reversal learning to improve reward outcomes (i.e., goal-directed action) over time, rather than a generalized increase in engagement with the operant device. An effect of psilocybin in early learning was not observed but this may be due to performance instability during this initial period (**Figure 5H-I**; *p*=.0657 and *p*<.05, respectively).

We have previously shown that psilocybin administered at this same dose did not affect effortful responding in rats using a progressive ratio of reinforcement as a measure of incentive motivation (27). However, to rule out possible contributions to task performance from general locomotor activity and anxiety-like behavior in the present study, we examined the effects of psilocybin on behavior in the OF (**Figure 5L**) and EPM (**Figure 5P**) in a separate cohort of animals. Psilocybin did not alter locomotor activity or change the duration spent in risky or safe zones of either the OF (**Figure 5M-O**) or EPM (**Figure 5Q-R**; all *ps* >.3217), indicating that these behavioral features did not explain the increase in task performance observed.

## Discussion

This study explored the enduring effects of psilocybin on behavior and information processing in rats, in an effort to further understand the mechanisms of how psilocybin may treat conditions such as depression. Our results support the notion that post-acute effects of psilocybin increase engagement in a reversal learning task, and suggest this may be explained by altered forgetting rates and reduced loss aversion. These results could have translational/clinical relevance to conditions, such as major depression, which are linked to a withdrawal from the world, and indeed existing (e.g., behavioral activation) treatments often specifically aim to promote greater engagement. Our results show the increased engagement could be due to increased optimism via an asymmetry in belief updating, highlighting optimism as a possible treatment target.

Our computational results show that psilocybin reduced belief updating rates after losses and increased belief updating rates after wins; that is, over time, the rats treated with psilocybin vs. controls forgot more about their previous beliefs when receiving a reward compared to when receiving a loss. These results are in line with an optimism bias, which relates to an asymmetry in belief updating in which individuals learn more from positive than negative outcomes (26,34,48,49). Interestingly, this biased belief updating manifests as more engagement in the environment. If an individual holds the (biased) belief that an action will lead to a positive outcome, they are more likely to choose that action – engaging more, and enabling themselves to minimize missed opportunities. Through this increased engagement with the world, optimism has been associated with improved quality of life and is suggested to be adaptive (26,50–58). As the psilocybin-treated rats increase the asymmetry in their belief updating, they also increase their engagement with the task. These results, along with complementary behavioral findings ruling out other explanations of increased engagement, support this optimism-based interpretation. Additionally, our results complement recent work showing that the antidepressant properties of ketamine treatment may be due to increased optimism from asymmetry in belief updating (59).

It is important to note that the rats given the saline control also showed asymmetry in their belief updating, where they had a lower forgetting rate for losses vs. rewards. This is not surprising, since research shows that some optimism in rats is widely present in healthy animals, and is absent mainly in rat models of depression (60,61). Our control group did not undergo any additional experimental interventions to develop pessimism, and would therefore be expected to show some optimism bias (62). Our findings suggest that psilocybin amplifies the asymmetry in updating seen in wild type rats. Future research on optimism should test the effects of psilocybin in a depression model cohort, such as in a model of chronic mild stress (63).

Our computational results also show that psilocybin reduced loss aversion in rats. The rats treated with psilocybin received more losses than the control group, but still engaged more with the task. As the rats had reduced loss aversion, they expected to dislike losses less. This means they were less deterred from engaging in the task than the control group, despite the possibility of a loss. As expected, this was also associated with more rewards. Our results thus suggest that, in addition to asymmetric belief updating, psilocybin may further increase engagement by reducing loss aversion. As loss aversion and pessimism are elevated in depression, these results are potentially promising in understanding how psilocybin may counter these crucial features of this disorder (22,34,64–66). Notably, these differences in loss aversion were seen in the later days of the task. This is consistent with human studies showing positive psilocybin effects weeks or months after treatment (67,68).

Previous studies on psilocybin and decision-making indicate that psilocybin may improve cognitive flexibility (69–71); a further study found that psilocybin may reduce punishment by changing one’s concerns for the outcome of their game partner in a social decision-making task (70). The current findings add to a growing literature in this area highlighting a range of effects that emerge in different task contexts, and which perhaps have similar underlying mechanisms. Additionally, our finding that changes in task engagement occur post-acutely complement research showing that after psilocybin treatment in mice, increases in dendritic spine peaked 7 days post-administration (72). The increased dendritic spine density may thus be required for the increased engagement.

Although it is beyond the scope of this paper to extend research on serotonin and psilocybin, our findings might also be relevant to existing theories on the role of serotonin. For example, one theory suggests that increased serotonin promotes exploration in order to identify new polices that can then be exploited (73). As female rats are expected to have increased serotonin (at least acutely) after treatment with psilocybin (27), and also explored their environment more, this could suggest a neural mechanism underlying our computational and other behavioral results. In our task increased engagement resulted in more rewards so the rat can then exploit increased engagement when serotonin levels reduce. This increased engagement policy results in optimistic learning of the task as the rat continues to receive reward. If serotonin does increase exploration after psilocybin treatment, it may have clinical implications when considering the environment within which a patient will explore and learn their new polices after treatment. In our task, performance also benefited from increased engagement; however, it should be noted that it is unknown at present whether this will generalize to other experimental or real-world contexts in which patients might benefit. Future studies should therefore extend our modeling approach to other environments and test potential links to underlying neurobiology.

Taken together, these results further support a way in which psilocybin treatment has potential to improve core symptoms of depression, associated with anhedonic, apathetic withdrawal, and diminished optimism, by altering specific computational mechanisms that lead to improved optimism and greater engagement with the world (14,15,34,64–66,74,75). Further research should consider if these effects on task engagement are specific to psilocybin, or if they can occur for other psychoactive agents (such as opiates), including how such psychoactive agents might manifest in the model parameters. This would be informative not only about the mechanism of psilocybin but also about different intervention targets; for example, even if opiates increase engagement, the underlying model might not reveal the same kind of belief updating asymmetry found here, which could have additional clinical relevance. Other relevant future research would include testing relevant antagonists and dose-dependence trials. It is also important to highlight that, as the clinical potential of these results depends on translatability to human participants, it will be crucial to find ways to ensure this is feasible. The current paradigm used here in rats required gathering behavior over a large number of sequential days, which could be challenging, especially in patient samples. Thus, adapted study procedures will likely need to be explored.

## Conclusion

In summary, we find that, in rats, psilocybin increases engagement with the environment consistent with an amplified optimism bias. Computational modelling suggests the psilocybin-treated rats have heightened expectations of reward as a result of changes in relative rates of belief updating from rewards and losses, as well as reduced loss aversion. This result has translational potential, and should motivate confirmation of these effects in human studies.

## Supplementary Materials

### Data availability

Code and data used presented in this paper are openly available on the Open Science Framework repository (DOI 10.17605/OSF.IO/EU8KH). Note: The original FED3 recordings did not include ‘Stay in Cage’ trials. The original data along with the code implemented to include ‘Stay in Cage’ trials, and processed data are available online.

### Model validation and comparison

When assessing the accuracy (percentage of trials where the model assigned the highest probability to the true actions) and average action probability (average probability the model assigned to the true actions) of each model, the 5-parameter AI model showed the best performance (see **Table S1**). Bayesian model comparison also confirmed this to be the winning model (protected exceedance probability = 1). For the winning model, recoverability analyses confirmed that generative and estimated parameters were significantly correlated: action precision (*r*=0.85, *p*<.001), forgetting rate for losses (*r*=0.57, *p*<.001), forgetting rate for rewards (*r*=0.94, *p*<.001), loss aversion (*r*=0.77, *p*<.001), and prior reward probability (*r*=0.84, *p*<.001). These results suggest that the winning model was reliable and accounted well for the behavioral data.

**Table S1.** Accuracy and average action probability of each model.

| Model | Accuracy | Average action probability |
| --- | --- | --- |
| RL simple | 32.76% | 0.37 |
| RL pessimism | 44.76% | 0.46 |
| RL associability | 21.06% | 0.33 |
| RL extended | 34.22% | 0.30 |
| AI 5 | 79.00% | 0.62 |
| AI 4 | 77.12% | 0.60 |

\*Note that chance accuracy is 33.33%

### Raw behavioral data

Supplementary Figure 1 shows the unsmoothed behavioral data. The mean for each group is shown for each day with standard error bars.

**Supplementary Figure 1:**
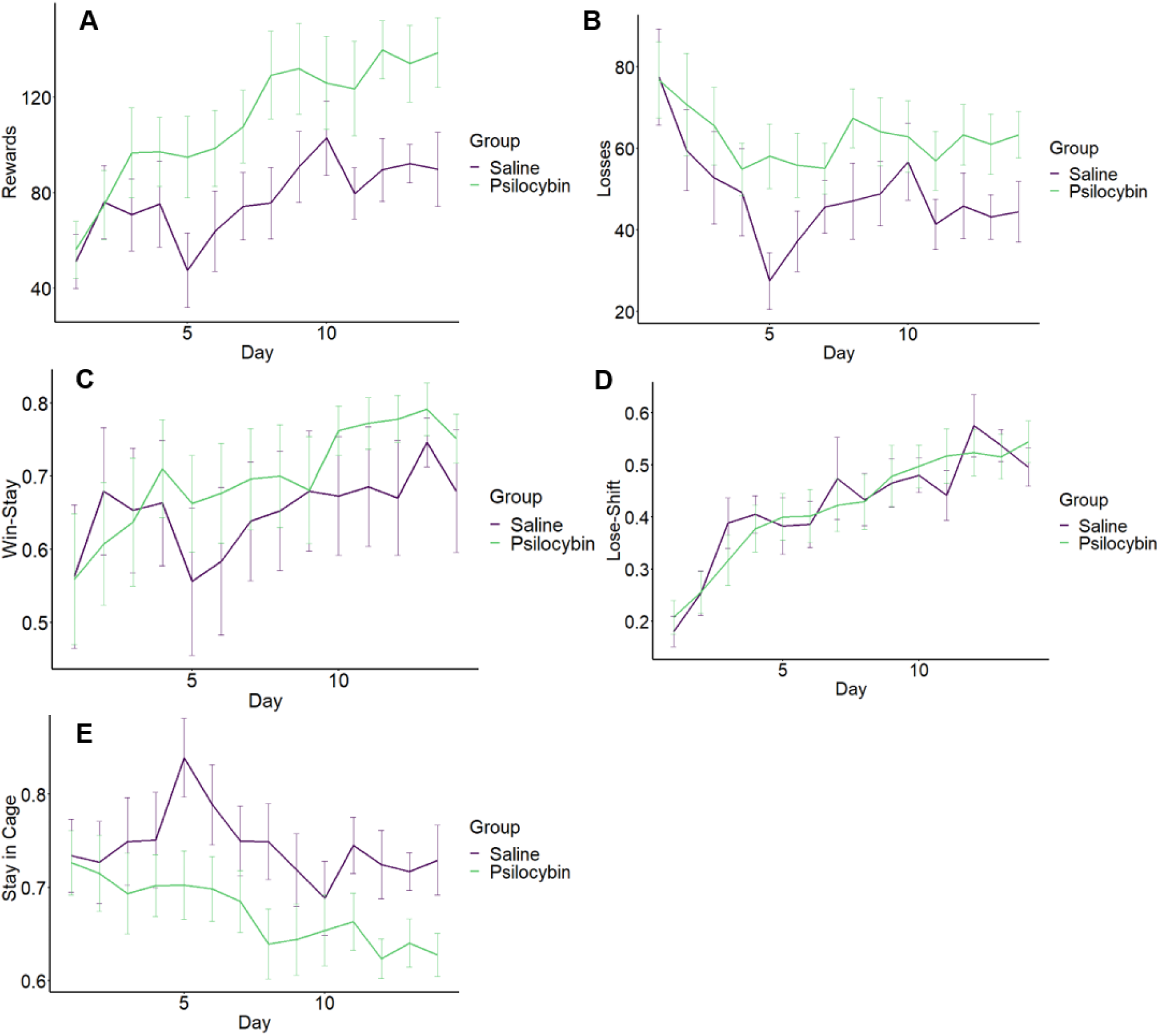
Effects of psilocybin on behavioral measures across days of reversal learning with unsmoothed data. The panels in this figure illustrate the mean of the behavioral measures with standard error bars across days including (A) rewards, (B) losses, (C) win-stay strategies, (D) lose-shift strategies and (E) stay in cage behavior.

### Raw model parameter data

Supplementary Figure 2 shows the unsmoothed model parameter data. The mean for each group is shown for each day with standard error bars.

**Supplementary Figure 2:**
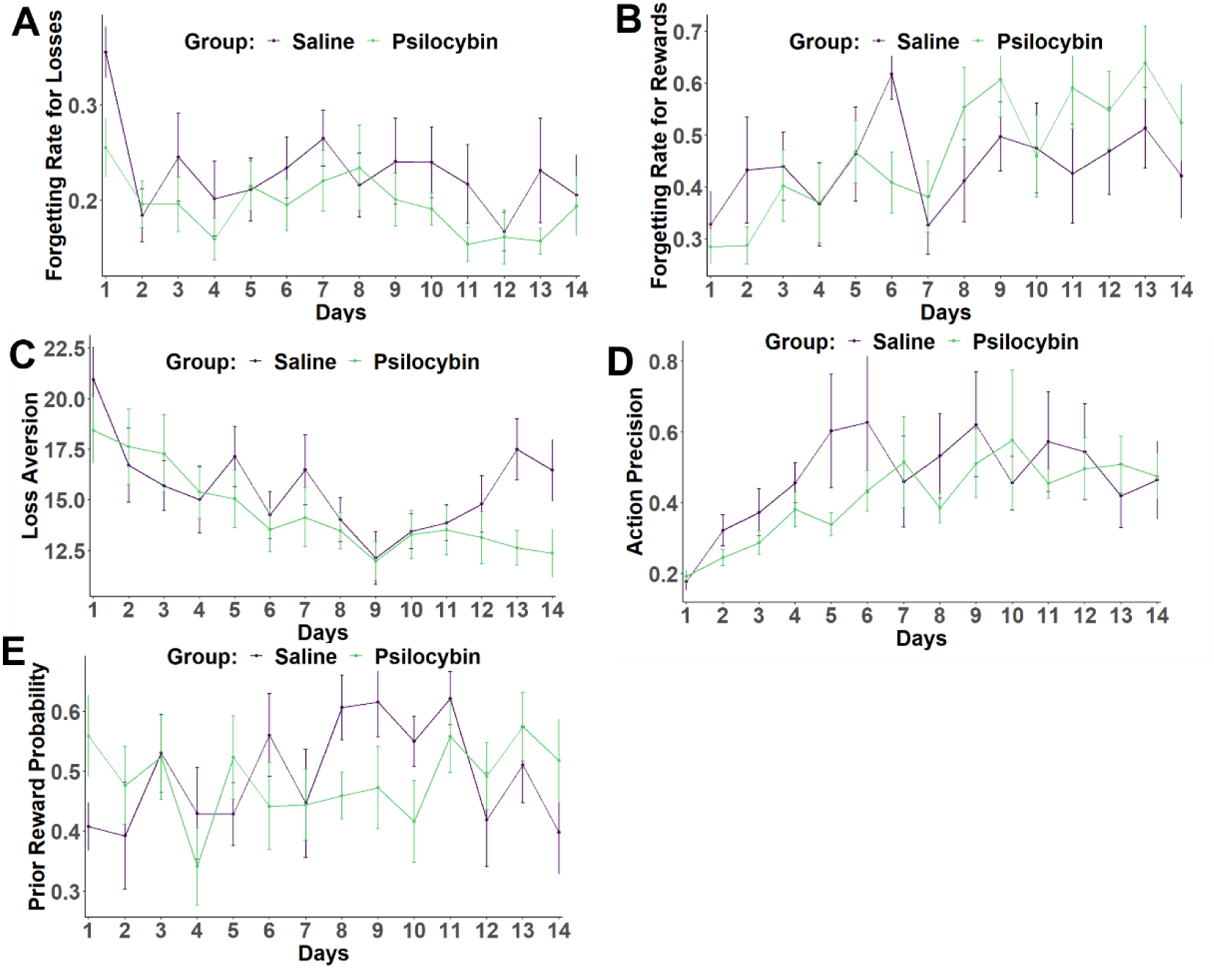
Effects of psilocybin on model parameters across days of reversal learning with unsmoothed data. The panels in this figure illustrate the mean of the model parameters with standard error bars across days including (A) forgetting rate for losses, (B) forgetting rate for rewards, (C) loss aversion, (D) action precision and (E) prior reward probability.

## Acknowledgments and Disclosures

This work was supported by a National Health and Medical Research Council Grant (Grant No. GNT2011334) to CJF and a Three Springs Foundation Grant awarded to JH. CJF sits on the scientific advisory board for Octarine Bio, Denmark. No other authors report biomedical financial interests or potential conflicts of interests

